# A type VII secretion system in Group B *Streptococcus* mediates cytotoxicity and virulence

**DOI:** 10.1101/2021.06.15.448449

**Authors:** BL Spencer, U Tak, JC Mendonça, PE Nagao, M Niederweis, KS Doran

**Affiliations:** University of Colorado Anschutz Medical Campus, Department of Immunology and Microbiology, Aurora, CO, USA; University of Alabama at Birmingham, Department of Microbiology, Birmingham, AL, USA; Rio de Janeiro State University, Roberto Alcântara Gomes Biology Institute, Rio de Janeiro, RJ, Brazil

**Author notes:** Present address: University of Colorado Boulder, Department of Biochemistry, Boulder, CO. **Correspondence:** Kelly S. Doran.

**Keywords:** *Streptococcus agalactiae*, Group B *Streptococcus*, meningitis, type VII secretion, pore formation, cytotoxicity, EsxA

## Abstract

Type VII secretion systems (T7SS) have been identified in Actinobacteria and Firmicutes and have been shown to secrete effector proteins with functions in virulence, host toxicity, or interbacterial killing in a few genera. Bioinformatic analysis indicates that Group B streptococcal (GBS) isolates encode four distinct subtypes of T7SS machinery, three of which encode adjacent putative T7SS effectors with WXG and LXG motifs. However, the function of T7SS in GBS pathogenesis is not known. Here we show that the most abundant GBS T7SS subtype is important for virulence and cytotoxicity in brain endothelium and that these phenotypes are dependent on the WXG100 effector EsxA. We further show that the WXG motif is required for cytotoxicity in brain endothelium and that EsxA is a pore-forming protein. This work reveals the importance of a T7SS in host–GBS interactions and has implications for the functions of T7SS effectors in other Gram-positive bacteria.

## Introduction

Bacteria utilize secretion systems to respond to changes in environment, defend against interbacterial killing, acquire nutrients, exchange genetic material, and promote virulence within the host ^1,2^. To date, several secretion systems have been identified in bacteria, but the majority are encoded by Gram-negative organisms. The type VII secretion system (T7SS) was discovered in *Mycobacterium tuberculosis* (*Mtb*), in which core machinery components assemble in the inner membrane and utilize an ATPAse to drive secretion of typically small, α-helical proteins lacking traditional signal peptides ^3^. These proteins are approximately 100 amino acids in length and center a tryptophan-variable-glycine (WXG) motif within the hairpin loop between two ɑ-helices; they are designated WXG100 proteins and are now considered canonically secreted factors of T7SSs ^4–6^. The five ESX systems in *Mtb* ^3^ secrete at least 22 WXG100 proteins ^7^ and have been implicated in a number of functions, including phagosomal rupture and macrophage intracellular survival ^8,9^, toxin secretion ^10^ and nutrient acquisition ^11^.

Improvements in next generation sequencing techniques have facilitated the identification of additional T7SS loci in other Actinobacteria (T7SSa) and in Firmicutes (T7SSb) based on homology of ATPase- and WXG100 protein-encoding genes ^7^. In Firmicutes, the mechanism and components of the T7SSb have been most extensively studied in *Staphylococcus aureus* ^12–16^, in which the core machinery consists of four membrane proteins: EsaA, EssA, EssB, and EssC, as well as a cytoplasmic protein, EsaB ^17,18^. While deletion of any one of these core components can abrogate T7SSb activity ^13,19,20^, the hexameric, membrane-bound ATPase EssC is considered the driver of T7SSb, of which the ATP-binding domains in the C-terminal region are required for substrate secretion and the C-terminal region is also necessary for substrate recognition and specificity ^15,21–23^. It has been shown in several Gram-positive bacterial species that the C-terminal sequence of EssC is highly variable across strains, resulting in EssC subtypes that are typically accompanied by a unique set of putative secreted effector-encoding genes ^17,24,25^. Thus, the effect of the T7SS on a bacterium’s interaction with other normal flora or on fitness within the host may be strain- or EssC subtype-specific.

Despite large variation in EssC subtypes and their cognate putative effectors between strains and bacterial species, genomic analyses indicate that T7SSb loci encode relatively conserved core components (including the N-terminus of EssC) as well as WXG100 protein EsxA, a widely studied T7SS substrate ^12,17,26,27^. Increasing numbers of reports have shown a role for the T7SSb and/or EsxA in the pathogenesis of several Gram-positive bacteria ^12,28–31^; however, T7SSb has not yet been characterized in the important pathogen *Streptococcus agalactiae* (also known as Group B *Streptococcus,* GBS). GBS is a β-hemolytic streptococcal species and the leading etiologic agent of bacterial meningitis in neonates ^32–34^. GBS exists primarily as an asymptomatic colonizer of the gastrointestinal and female reproductive tracts but can cause disease in newborns upon its transmission from the vaginal tract of the mother *in utero* or during birth ^35,36^. In the newborn, GBS can infect the lungs or blood to cause pneumonia and bacteremia, and in some cases may penetrate the brain resulting in meningitis ^37,38^. GBS infection among other immunocompromised populations such as elderly adults or adults with cancer or diabetes is also rising in prevalence ^39–41^. While many factors have been identified that mediate the physical interaction of GBS with the brain endothelial cells that constitute the blood-brain barrier (BBB) ^42^, the mechanisms by which GBS damages or breaks down that endothelial barrier are still being elucidated.

Herein, we characterize the T7SSb in GBS. We show by genomic analysis of available whole genome sequences that the GBS T7SS can be divided into at least four subtypes based on the C-terminus of EssC. The GBS T7SS subtype I is the most prevalent, representing >50% of all isolates analyzed. Using a representative subtype I GBS strain, CJB111, we show that deletion of the ATPase-encoding gene, *essC,* mitigates virulence and GBS-induced inflammation in the brain, as well as cell death in brain endothelial cells and that these phenotypes are dependent on the T7SS canonical substrate EsxA. We further show that the EsxA WXG motif is required for cytotoxicity in brain endothelium and that EsxA is a pore-forming protein. Our study provides the first experimental evidence indicating the T7SS promotes GBS pathogenesis and is the first demonstrate a role for a non-mycobacterial EsxA homolog in pore formation.

## Results

### Identification of four GBS T7SS subtypes based on EssC protein sequence

As a T7SS for major neonatal pathogen GBS has not been described, we analyzed closed genome sequences from GBS isolates for the presence of T7SS core genes and putative effectors. We observed an extensive amount of genetic diversity in T7SS operons in regard to sequence homology of *essC*, the presence of one or two *esxA* homologs, as well as the presence of putative T7SS effectors and putative LXG toxin/anti-toxin-encoding genes downstream of the core T7SS machinery genes. To determine which GBS T7SS subtype might be most prevalent, we examined the C-terminal 225 amino acids of EssC. In *S. aureus* and *Listeria monocytogenes,* the EssC C-terminus is the point at which the protein sequence diverges into distinct EssC variants, and each associate with unique downstream putative effector-encoding genes ^15,25^. Of the 80 GBS whole genome sequenced isolates that encode the 225 C-terminal amino acids of EssC (Table S1), the majority (46/80; 57.5%) encode an EssC variant that we now classify as subtype I (Fig. 1a-b).

**Fig. 1.**
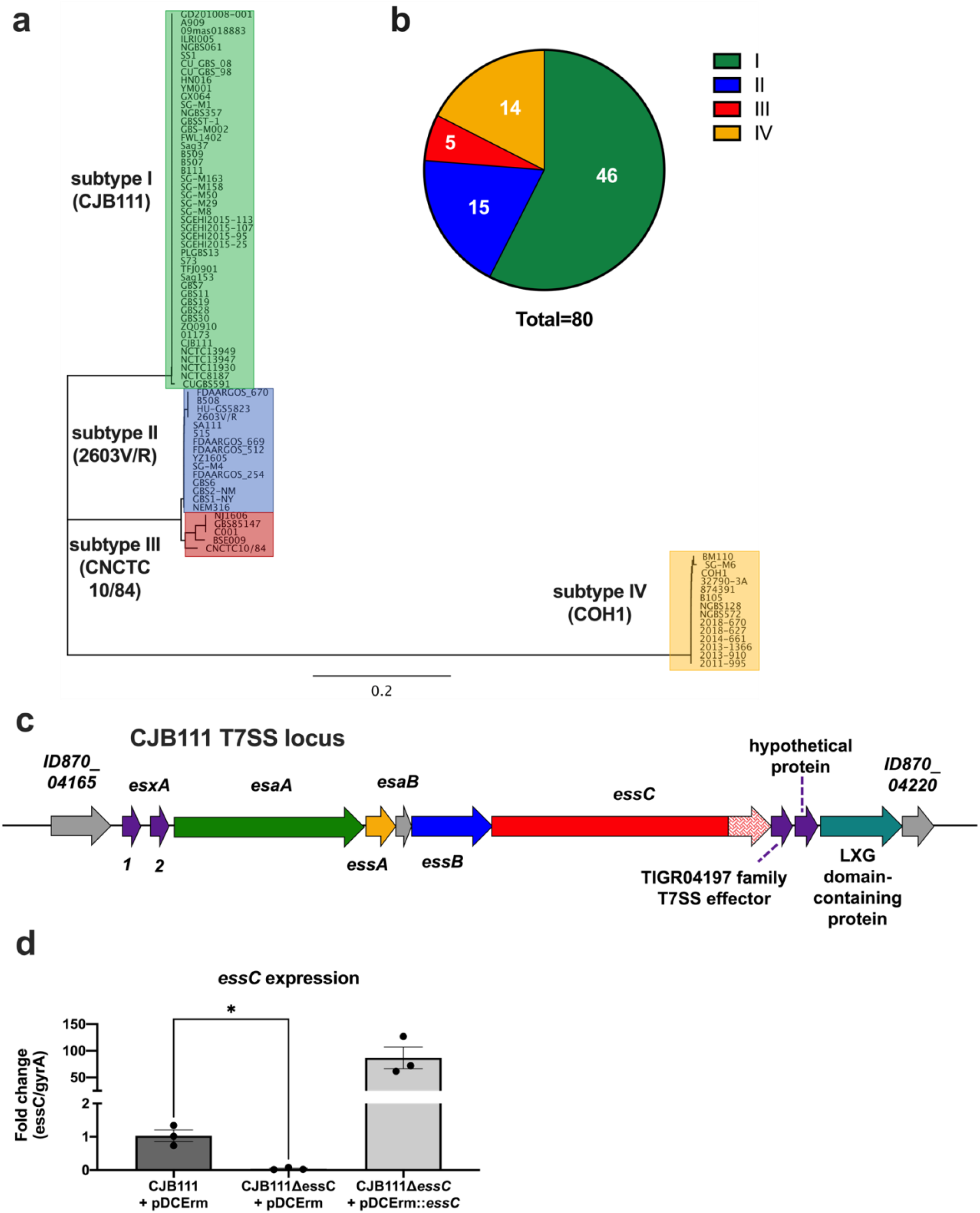
Genomic analysis of GBS T7SS subtypes and characterization of the GBS T7SS operon in subtype I strain, CJB111. **a)** Phylogenetic tree of whole-genome sequenced GenBank isolates that encode the C-terminal 225 amino acids of EssC. Subtype I strains (represented by strain CJB111; n = 46) are highlighted in green, subtype II strains (represented by 2603V/R; n = 15) are highlighted in blue, subtype III strains (represented by CNCTC 10/84; n = 5) are highlighted in red, and subtype IV strains (represented by COH1; n = 14) are highlighted in yellow. **b)** Distribution of GBS T7SS subtypes based on EssC C-terminus in whole-genome sequenced GBS isolates that encode T7SS (n = 80). **c)** Diagram of the T7SS locus in CJB111 (roughly to scale; accession CP063198.2). Genes in purple and teal encode WXG100 proteins or putative T7SS secreted effectors. Putative core genes of the operon are *esaA* through *essC*. **d)** Expression of *essC* in CJB111+ pDC, CJB111Δ*essC*+ pDC, CJB111Δ*essC* + pDC*essC* strains by qRT-PCR. T7SS gene expression was normalized to housekeeping gene *gyrA.* Statistics reflect the repeated measures, one way ANOVA with Dunnett’s multiple comparisons test to CJB111Δ*essC*+pDC, *p* < 0.05, *. Data indicate the mean of three independent experiments and error bars represent standard error of the mean.

### Characterization of GBSS T7SS in subtype I strain, CJB111

In this study, we have utilized neonatal isolate CJB111 as a representative subtype I strain to study the role of the GBS T7SSb in virulence. In addition to EssC, CJB111 encodes T7SSb core components EsaA, EssA, EssB, and EsaB, which are homologous to those found in *S. aureus* genomes (Fig. 1c). CJB111 also encodes two copies of the WXG100 protein-encoding gene *esxA* (designated as *esxA1* and *esxA2*) upstream of the T7SS core genes and additional putative T7SS effectors (including an LXG-domain containing protein) downstream of the T7SS core genes (Fig. 1c). In *S. aureus*, the EssC ATPase has been characterized as the driver of T7SS secretion and deletion of this gene abrogates secretion of all T7SS substrates ^12,19^; we hypothesized that EssC might have a similar role in GBS and thus deleted *essC* from CJB111 to assess the contribution of T7SS to GBS pathogenesis. The Δ*essC* deletion strain was complemented using an overexpression plasmid and these strains were confirmed to have the expected expression levels of *essC* (Fig. 1d).

### T7SS contributes to virulence and meningitis development

To assess if the GBS T7SS is important for virulence *in vivo*, we infected CD1 mice with CJB111 or CJB111Δ*essC* via tail vein injection and assessed survival and meningitis progression. Mice displaying neurological symptoms, such as paralysis or seizures, as well as those found moribund were sacrificed to generate a survival curve between the two groups. Mice injected with CJB111 succumbed to infection much more quickly than those infected with the Δ*essC* mutant. By 36 hours post-infection, approximately 55% of the CJB111-infected mice had succumbed to infection compared to just 5% of those infected with the Δ*essC* mutant (Log rank test, *p* = 0.0001, ***; Fig. 2a). To assess the impact of the T7SS on GBS burden during infection, blood and various tissues were isolated, homogenized and plated to enumerate CFU (Fig. 2b-d). Significantly less bacteria were observed in the blood and heart of Δ*essC-*infected mice (Fig. 2b, c) implicating CJB111 T7SS in virulence of GBS. However, we did not observe a significant effect of the T7SS on bacterial burden in the brain (Fig. 2d). This is consistent with our finding that the Δ*essC* mutant exhibited a similar ability to adhere to, invade, and survive in human cerebral microvascular endothelial cells (hCMEC), which we used as an *in vitro* model for the BBB ^43^ (Fig. S1a-c). Because meningitis is an inflammatory disease, we hypothesized that CJB111 might elicit a heightened inflammatory response in the brains of infected mice, despite the relatively equivalent bacterial load observed in CJB111- and Δ*essC* mutant-infected mice. Using ELISA to quantify protein levels in brain tissue, we observed that mice infected with CJB111 had significantly higher amounts of the neutrophil chemokine KC (an early indicator of meningitis development) in brain tissue than those infected with the Δ*essC* mutant (Fig. 2e). Further, when KC protein levels were normalized to the bacterial CFU within the same tissue, brains of CJB111-infected mice exhibited higher levels of KC protein compared to brains of *essC* mutant-infected mice (Fig. 2f). This demonstrates that even at equivalent bacterial tissue burden, T7SS-sufficient CJB111 elicits more inflammation in the brain compared to the Δ*essC* mutant, which may promote the large survival differences observed between groups during meningitis progression.

**Fig. 2.**
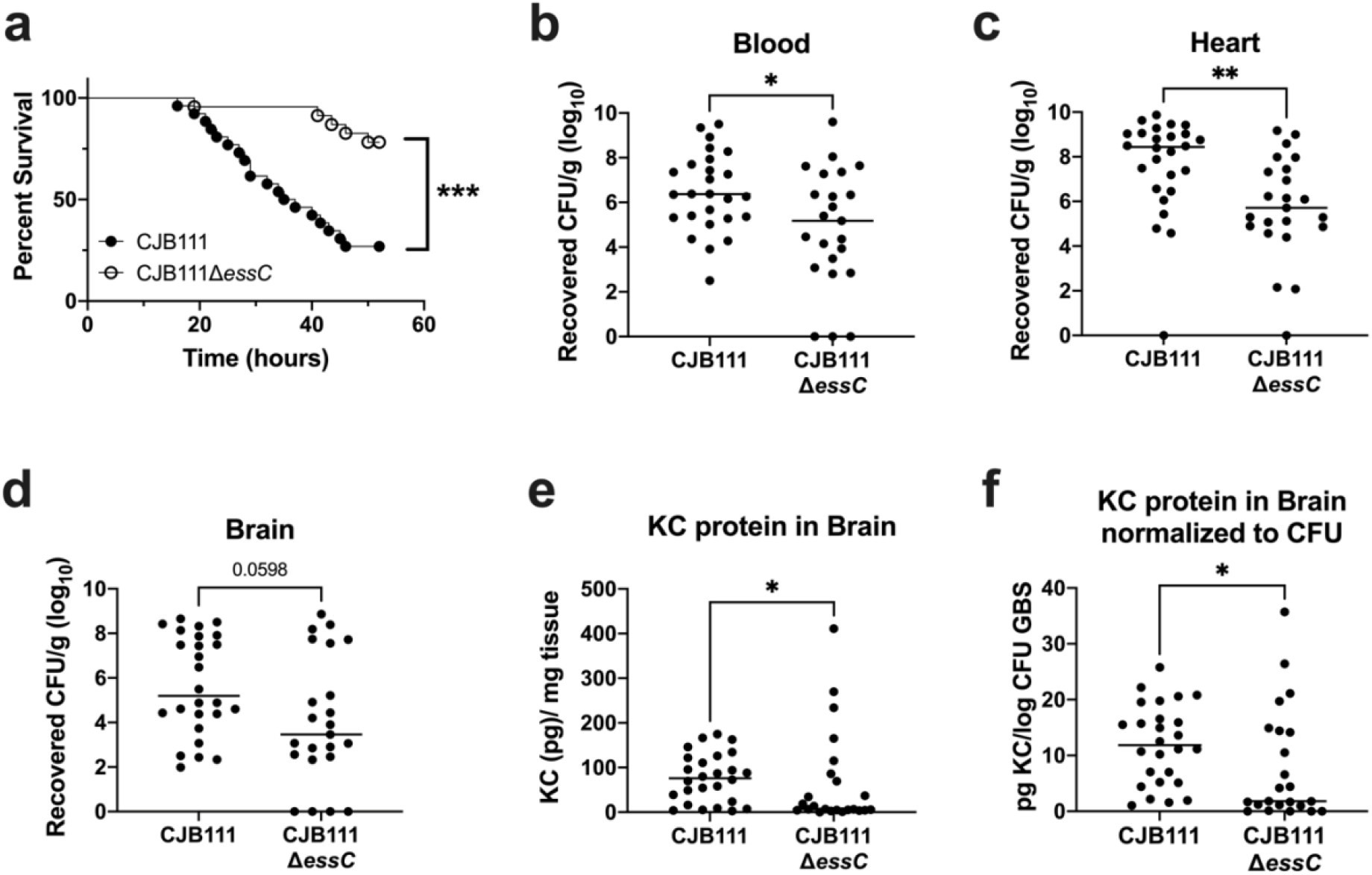
CJB111 T7SS mediates virulence in a model of hematogenous meningitis. **a)** Survival curve of 8 week-old CD1 male mice tail-vein injected with CJB111 (n = 26) or CJB111Δ*essC* (n = 23). Graph shows combined survival curves of three independent experiments, all of which ended at 52 hours post-infection. Statistics reflect the Log rank (Mantel-Cox) test, *p* < 0.001, ***. Recovered CFU counts from the **b)** blood **c)** heart, and **d)** brain tissue of infected mice. **e)** KC protein levels quantified from infected brain tissue by ELISA, and **f)** normalized to the log_10_ CFU within each brain. In panels **b-f**, each dot represents one mouse and plots show the median. Statistics represent the Mann Whitney *U* test. *p* < 0.05, *; *p* < 0.01, **.

### CJB111 T7SS induces cell death in brain endothelial cells

Damage to host endothelium occurs during bacterial infection and sepsis ^44,45^ and can exacerbate disease progression and result in multi-organ failure ^46^. T7SS in other organisms, such as *S. aureus,* is known to secrete toxins that target the host ^16^. To determine if GBS T7SS induces endothelial cell death, we measured cytotoxicity induced by CJB111, CJB111Δ*essC* mutant, and complemented strains in hCMEC using lactate dehydrogenase (LDH) release assays (see Materials and Methods). CJB111 induced approximately 70% cytotoxicity after an infection of MOI 10 for 4-5 hours, while cytotoxicity caused by the Δ*essC* mutant was significantly reduced (~40%). This phenotype was complemented by expression of *essC* in the Δ*essC* mutant (Fig. 3a). This T7SS-mediated cytotoxicity was largely contact-dependent as experiments in which the bacteria and cells were separated by a 0.4 μm transwell resulted in minimal LDH release (~4% cytotoxicity), approximately 15 times lower than the cytotoxicity observed during normal infection by CJB111 (Fig. 3b). However, even this slight level of contact-independent cytotoxicity was T7SS-dependent as the Δ*essC* mutant did not induce detectable levels of cytotoxicity compared to untreated cells (Fig. 3b). These data indicate that the CJB111 T7SS mediates cell death responses in brain endothelium that may translate to poor outcomes during *in vivo* infection models.

**Fig. 3.**
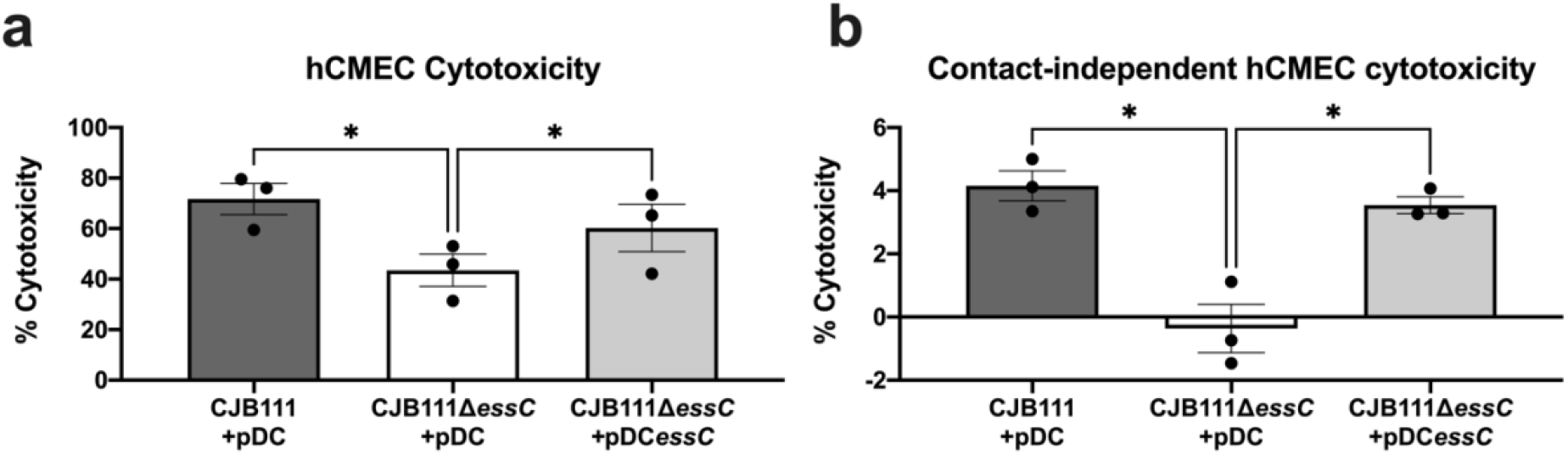
CJB111 T7SS elicits cell death in human cerebral microvascular endothelial cells. **a)** hCMEC cytotoxicity calculated based on LDH release assay. Supernatant was collected from hCMECs infected with CJB111+pDC, CJB111Δ*essC*+pDC, and CJB111Δ*essC*+pDC*essC* at MOI 10, 4-5 hours post-infection. Percent cytotoxicity was calculated based on 0% (uninfected) and 100% (+lysis buffer) lysis controls. **b)** Contact-independent cytotoxicity in hCMECs during GBS infection using 0.4 μm polycarbonate transwells in which GBS and cells were separated by the porous membrane. Supernatant from hCMEC compartments were plated after the experiment to ensure no bacterial contamination. Cytotoxicity was measured 24 hours later. Statistics reflect the repeated-measures one way ANOVA with Dunnett’s multiple comparisons to CJB111Δ*essC*+pDC, *p* < 0.05, *. Data represent the mean of three independent experiments and error bars represent standard error of the mean.

### CJB111 WXG100 protein EsxA homologs in other T7SS-encoding bacteria and other GBS isolates

The most conserved T7SS effector described in the literature is ESAT-6 (early secreted antigenic target of 6 kDa; also known as EsxA), which was the first identified secreted substrate of the *Mtb* T7SS ESX-1 ^47,48^. EsxA orthologs are encoded across Actinobacteria and Firmicutes and retain a similar antiparallel hairpin structure despite large differences in sequence identity ^4^. GBS strain CJB111 encodes two adjacent EsxA homologs that are 95% identical to each other and are located immediately upstream of the T7SS locus (Fig. 1c). We performed BLAST analysis to compare the CJB111 EsxA1 against orthologs in other species in which the T7SS has been studied: *Mtb* (H37Rv) ^47^, *S. aureus* (USA300 strain FSRP357) ^13^, *Enterococcus faecalis* (OG1RF) ^29^, *L. monocytogenes* (EGD-e) ^49^, *Bacillus subtilis* (PY79) ^50^, *Streptococcus intermedius* (B196) ^30^*, Streptococcus suis* (GZ0565) ^51^, and *Streptococcus gallolyticus* (TX20005) ^31^. CJB111 EsxA1 shared the least identity with *Mtb* EsxA (13%) and was more similar to *S. aureus* EsxA (49% identical) than to *S. intermedius* or *E. faecalis* EsxAs (33% and 32% identical, respectively), despite the fact that *Streptococcus* and *Enterococcus* are more phylogenetically similar overall ^52^. Unsurprisingly, CJB111 EsxA1 displayed high identity with EsxA expressed by streptococcal species *S. suis* and *S. gallolyticus* (65 and 66% identical, respectively) (Fig. 4).

**Fig. 4.**
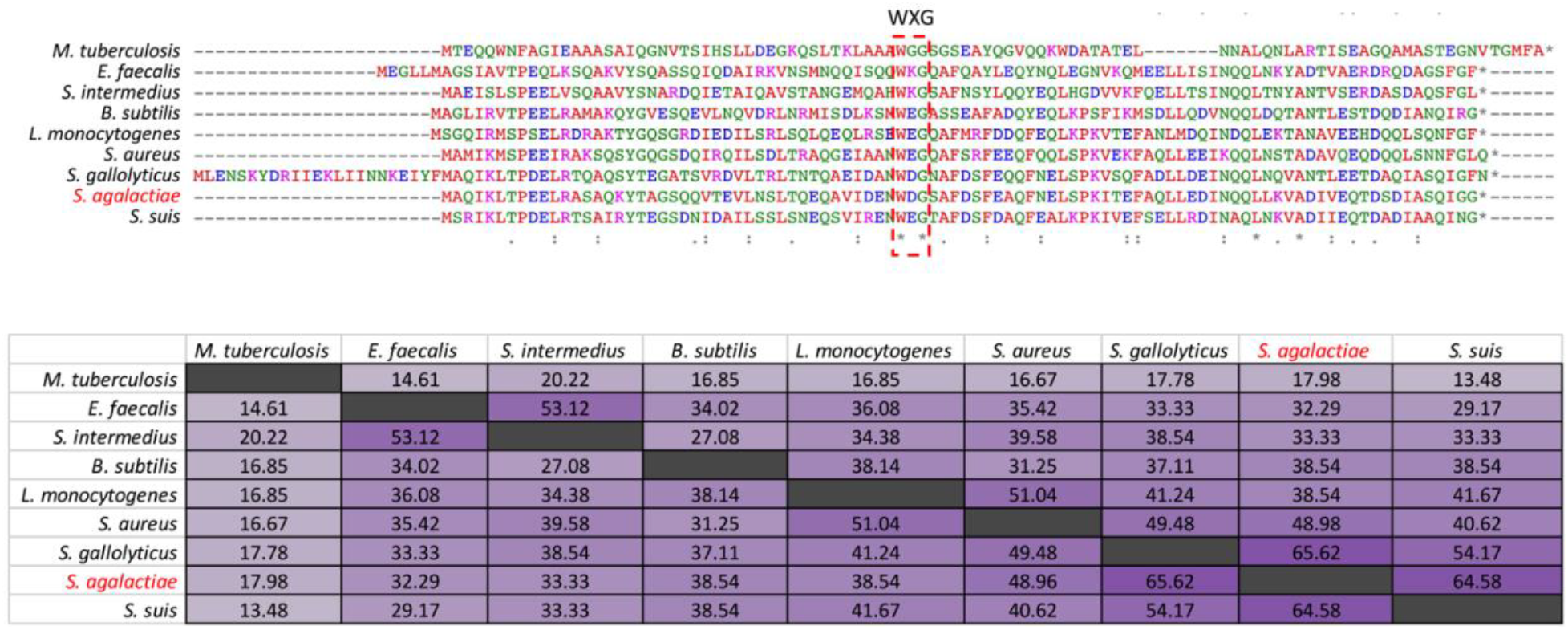
Canonical T7SS substrate EsxA is conserved across T7SSa and T7SSb loci. ClustalW alignments and percent identity matrix of EsxA protein sequences from *Mtb* (H37Rv; accession CP003248.2), *E. faecalis* (OG1RF; accession CP002621.1), *S. intermedius* (B196; accession NC_022246.1), *B. subtilis* (PY79; accession NC_022898.1), *L. monocytogenes* (EGD-e; accession NZ_CP023861.1), *S. aureus* (USA300 FPR3757; accession NC_007793.1), *S. gallolyticus* (TX20005; accession AEEM00000000.1), *S. agalactiae* (CJB111; accession CP063198.2) and *S. suis* (GZ0565; accession NZ_CP017142.1). In the above matrix, the purple shading corresponds to the level of identity between two strains (on a spectrum of 0 to 100% identity), with darker shading indicative of higher percent identity.

Within GBS, four T7SS subtypes exist based on the EssC C-terminal amino acid sequence and on presence/number of EsxA homologs. Similar to subtype I strain CJB111 (described above), subtype II GBS strains (represented by strain 2603V/R or NEM316) and subtype III GBS strains (represented by CNCTC 10/84) also encode EsxA upstream of the T7SS core genes. However, subtype II strains encode just one EsxA, which is most similar to CJB111 EsxA2 (98.98% identity; Fig. S2). Subtype III strains encode both EsxA1 and EsxA2, but EsxA2 is truncated due to a frameshift approximately 75% through the coding sequence. It is unknown whether this truncated EsxA2 remains functional. Finally, subtype IV, represented by strain COH1 (and other ST-17/serotype III GBS isolates), does not encode EsxA or any predicated WXG100 protein. As all characterized T7SS systems to this point have contained both an FtsK/SpoIII ATPase and a WXG100 homolog, it is unclear what the function of T7SS in subtype IV GBS strains might be.

### EsxA contributes to CJB111 virulence and cytotoxicity

To determine if EsxA contributes to T7SS-dependent virulence *in vivo*, we constructed a CJB111 mutant lacking both *esxA1* and *esxA2* genes (referred to here as Δ*esxA1-2)*. CD1 mice were infected as described above and in Materials and Methods with parental CJB111 and Δ*esxA1-2* mutant strains and monitored for mortality and moribundity. We observed that mice infected with the Δ*esxA1-2* mutant exhibited no mortality compared with the 75% mortality that occurred in mice infected with the parental CJB111 strain (Log rank test, *p* = 0.0013, **; Fig. 5a). Further, mice infected with the Δ*esxA1-2* mutant had significantly lower bacterial burden in the blood, as well as in heart and brain tissue (Fig. 5b-d) compared to CJB111-infected mice. As observed previously in mice infected with the Δ*essC* mutant (Fig. 2e-f), there were decreased levels of neutrophil chemokine KC in the brain in Δ*esxA1-2* mutant-infected mice compared to CJB111-infected mice (Fig. 5e). This difference was also exacerbated upon normalization of brain KC protein levels to bacterial brain CFU (Fig. 5f). These data indicate a role for EsxA in both GBS virulence and meningitis progression.

**Fig. 5.**
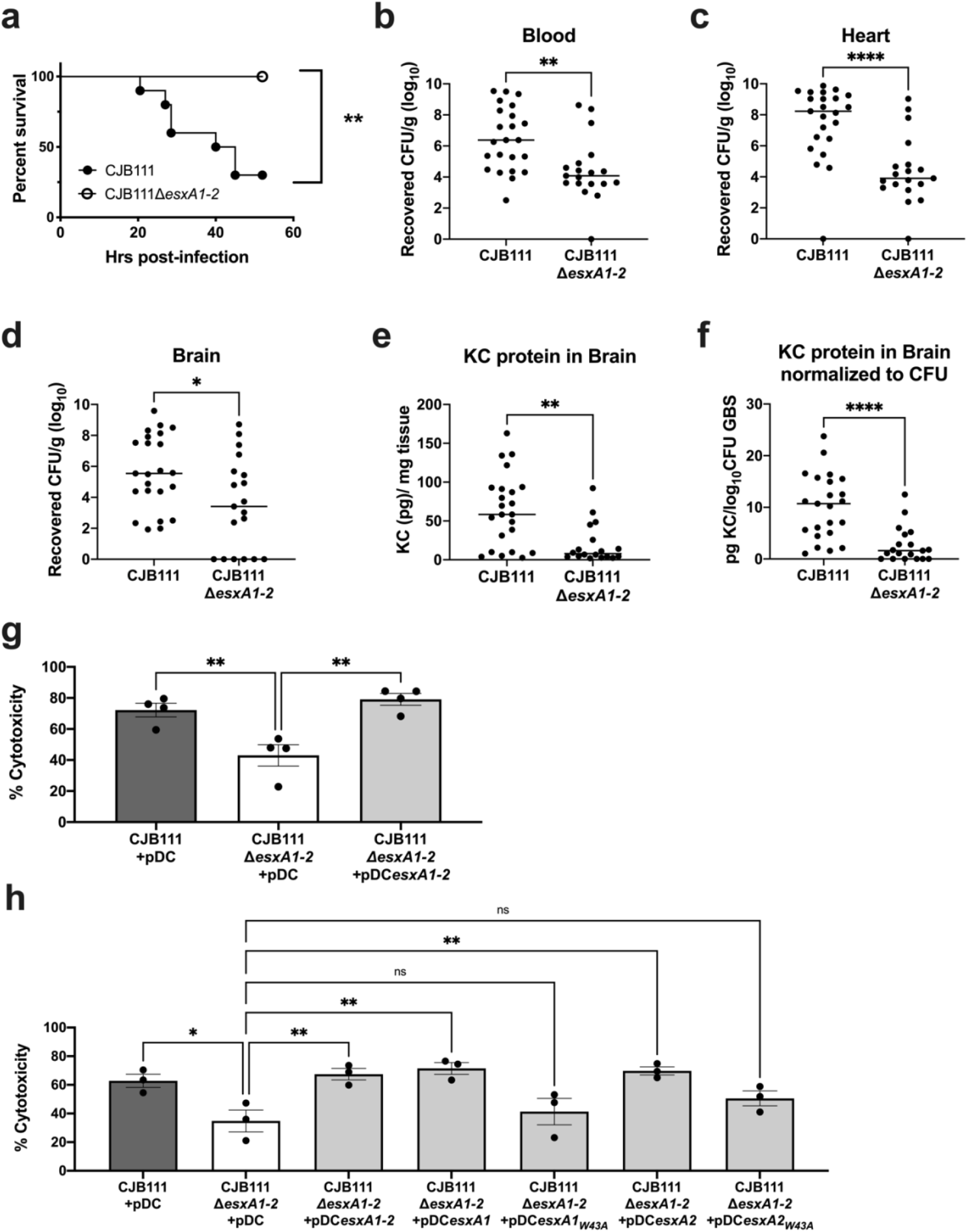
EsxA contributes to CJB111 virulence and cytotoxicity. **a)** Representative survival curve of 8 week-old CD1 male mice tail-vein injected with CJB111 or CJB111Δ*esxA1-2.* Graph shows survival curve of one representative experiment. n =10 mice/ group. Statistics reflect the Log rank (Mantel-Cox test), *p* < 0.01, **. Recovered CFU counts from the **b)** blood and **c)** heart, and **d)** brain tissue of infected mice. **e)** KC protein levels quantified from infected brain tissue by ELISA, and **f)** normalized to the log_10_ CFU within each brain. In panels **b-f**, each dot represents one mouse and plots show the median. Statistics represent the Mann Whitney *U* test. *p* < 0.05, *; *p* < 0.01, **; *p* < 0.0001, ****. **g-h)** Percent cytotoxicity calculated based on LDH release assay. In panel **g**), supernatant was collected from hCMECs infected with CJB111 + pDC, CJB111Δ*esxA1-2* + pDC, and CJB111Δ*esxA1-2* **+** pDC*esxA1-2* and in panel **h**), supernatant was collected from hCMECs infected with CJB111 + pDC, CJB111Δ*esxA1-2* + pDC, CJB111Δ*esxA1-2* **+** pDC*esxA1-2,* and single *esxA* complements CJB111Δ*esxA1-2* **+** pDC*esxA1*, CJB111Δ*esxA1-2* **+** pDC*esxA1_W43A_*, CJB111Δ*esxA1-2* **+** pDC*esxA2*, and CJB111Δ*esxA1-2* **+** pDC*esxA2_W43A_*, at MOI 10, 4-5 hours post-infection. Percent cytotoxicity was calculated based on a 0% (uninfected) and 100% (+lysis buffer) controls. In panels **g-h**, statistics reflect one way ANOVA with Dunnett’s multiple comparisons to CJB111Δ*esxA*1-2 + pDC, *p* < 0.05, *; *p* < 0.01, **. Data represent the mean of at least three independent experiments and error bars represent standard error of the mean.

Finally, to determine if EsxA is contributes to the T7SS-dependent cytotoxicity phenotypes observed in brain endothelial cells (Fig. 3), hCMEC monolayers were infected with CJB111, Δ*esxA1-2* mutant, and complemented strains. Similar to the Δ*essC* mutant, the Δ*esxA1-2* mutant exhibited attenuated cytotoxicity in brain endothelium compared to the parental CJB111 strain (Fig. 5g). This EsxA-dependent cytotoxicity in hCMECs could be restored with a double *esxA1esxA2* complement or with *esxA1* or *esxA2* single complements (Fig. 5h), indicating that expression of either of the EsxA proteins is sufficient for T7SS-dependent cytotoxicity.

WXG100 proteins such as EsxA form antiparallel ɑ-helical bundles, with the hydrophobic WXG motif located in the hairpin loop ^4^. These WXG motifs have been shown to facilitate oligomerization and pore formation and may also mediate export of other T7SS substrates ^10,27,53^. To assess the influence of the EsxA WXG motif on host cell cytotoxicity, we created single gene complements expressing WXG-mutant EsxA1 or EsxA2 (annotated as *esxA1*_W43A_ or *esxA2*_W43A_); yet, neither of these significantly complemented the cytotoxicity defect of the Δ*esxA1-2* mutant (Fig. 5h). These data indicate that the WXG motif is indeed important for EsxA-mediated cytotoxicity in brain endothelium.

### CJB111 EsxA1 is a pore-forming protein

EsxA is a well-known T7SS substrate ^3,12,20,26,54^; yet the specific mechanism by which it contributes to T7SS-dependent phenotypes is not clearly defined. Tak et al. recently showed that the EsxE-EsxF complex, a mycobacterial WXG100 protein pair, forms pores to enable toxin secretion ^10^. Thus, we hypothesized that GBS EsxA might also form pores, facilitating our observed T7SS-dependent phenotypes. To examine this, we purified CJB111 EsxA1 as described in the Methods via expression of an EsxA1-maltose binding protein (MBP) fusion ^10,55^ (Fig. S3a-c). EsxA1 was purified in the absence of detergents to prevent potential artifacts ^56^. Similar to previous purifications of Esx proteins ^10^, GBS EsxA1 formed many oligomeric species, which were confirmed by native PAGE using EsxA1 antiserum (Fig. S3d). Most of the high molecular weight oligomers were dissociated to the monomer by 6 M guanidine hydrochloride, except for a protein species with an apparent molecular weight of approximately 100 kDa. This band stained with anti-EsxA1 antiserum, but not with an anti-MBP antibody (Fig. S3d), indicating it is a stable oligomer of EsxA1.

To determine whether GBS EsxA1 forms pores, we used planar lipid bilayer experiments as previously described ^10^. While we observed no channel activity with buffer alone, the purified GBS EsxA1 protein formed transmembrane pores, as observed by a stepwise current increase (Fig. 6a-b, Fig. S4). Interestingly, reducing the buffer pH from 7.4 to pH 4.0 increased the channel activity significantly (Fig. 6b, Fig. S4) demonstrating that EsxA1 forms open pores in lipid membranes. A strong pH dependency of the channel-forming activity was also observed for the EsxE-EsxF complex with a 10-fold reduced channel activity at pH 5.5 compared to pH 6.5 ^10^.

**Fig. 6.**
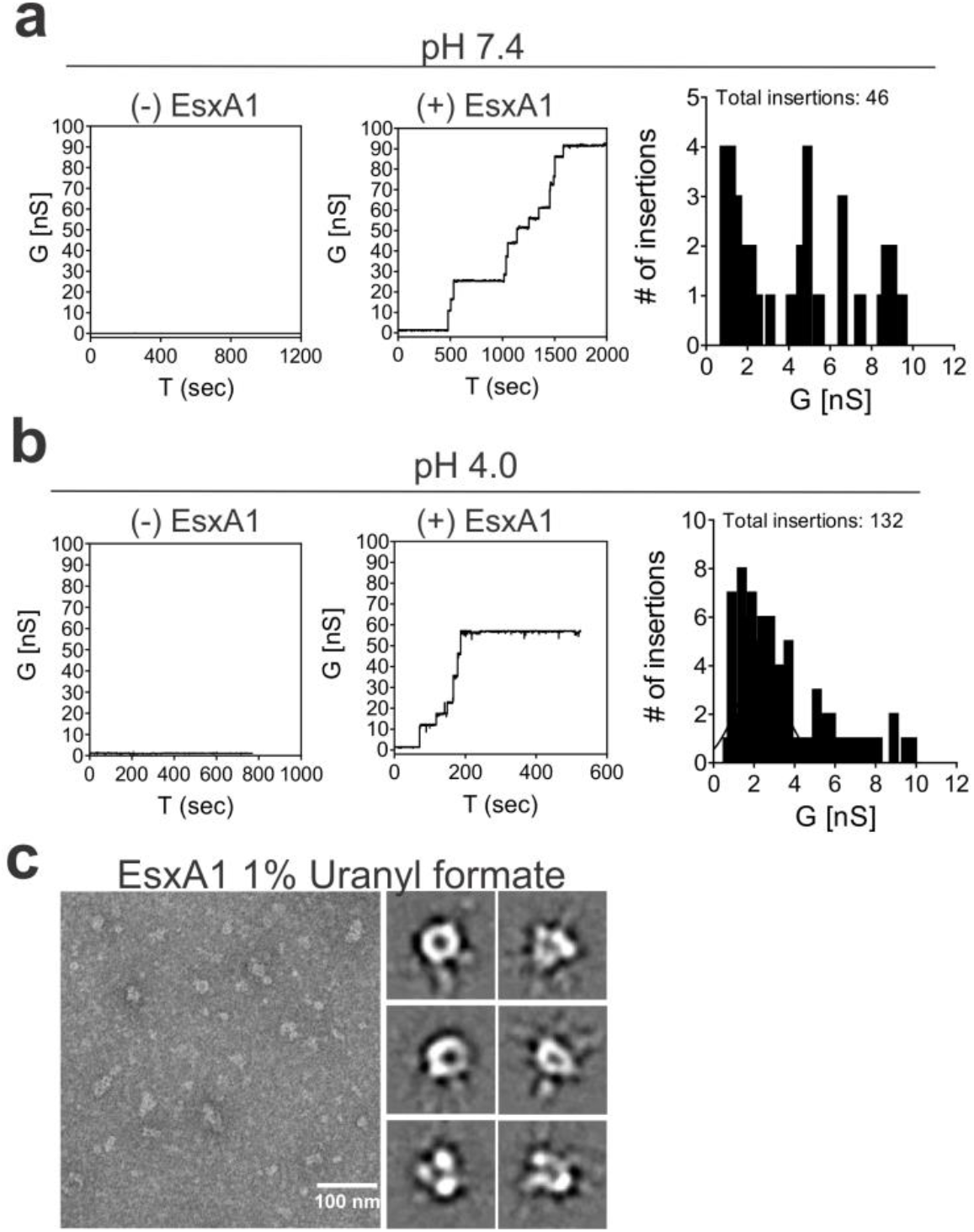
CJB111 EsxA is a pore-forming protein. Lipid bilayers composed of 1,2-diphytanoyl-sn-glycerol-3-phosphocholine (DphpC), 4ME 16:0 PC were incubated with 5 μg CJB111 recombinant EsxA at **a)** pH 7 and **b)** pH 4.0 in 25 mM sodium phosphate 1M KCl. Representative current traces are shown, and insertion sizes and frequencies are summarized in the histograms. Ten membranes were run in each buffer condition. No protein controls were also performed to rule out artifactual pore formation due to contamination of the system or buffer. Additional current traces can be found in Fig. S4a-b. **c)** Transmission electron microscopy of recombinant EsxA1. Shown are EsxA1 particles negatively stained with 1% uranyl formate and selected reference-free 2D class averages from 12,727 particles that resembled an intact oligomeric pore. The full set of class averages can be found in Fig. S4c.

Electron microscopy of negatively stained protein samples enables visualization of open channel proteins ^10,57^. Indeed, purified water-soluble EsxA1 protein negatively stained with uranyl formate and imaged by electron microscopy revealed particles with typical appearance of pores (Fig. 6c). Reference-free 2D class averaging of 12,727 particles revealed consistent oligomeric complexes with a central pore as well as multiple heterogeneous complexes that may represent incomplete or assembling pores (Fig. 6c, Fig. S4) thus corroborating our observations of higher-order oligomers by native gel (Fig. S3). Collectively, these results indicate that GBS EsxA1 forms water-soluble pore and/or pre-pore complexes that are capable of membrane insertion.

## Discussion

In this study, we describe the first role for the T7SS in GBS and identify four T7SS subtypes based on the C-terminus of the ATPase EssC. We further demonstrate a role for GBS T7SS subtype I in virulence and meningitis progression and show that it is dependent on conserved T7SS effector EsxA. Finally, we show that T7SS-and EsxA-dependent virulence in CJB111 may be promoted by the ability of EsxA to form pores in membranes and to induce cytotoxicity in brain endothelial cells via the canonical WXG motif.

As the most broadly conserved T7SS effector, EsxA is known to contribute to numerous virulence phenotypes in *Mtb* and, more recently, in Firmicutes ^54^. In *Mtb*, ESAT-6, or EsxA, was shown to promote virulence in a murine model of infection ^27^ and the ESX1 system that secretes EsxA has been well established as necessary for phagolyosomal escape and intracellular survival in macrophages ^8,9^. *Mtb* EsxA is also known to elicit strong interferon responses from T cells ^47^, is strongly immunodominant in the T-cell response to *Mtb* ^58,59^, and induces apoptosis and membrane perturbation in host cells ^60–62^. Similarly, in *S. aureus*, EsxA is important for virulence in models of infection, specifically in abscesses, and *S. aureus* T7SS has broadly been attributed to virulence in murine models of blood infection, nasal colonization, and pneumonia ^12,19,63,64^.

While it is clear that the T7SS plays a role in virulence in several bacterial species, the specific mechanism by which EsxA or other WXG100 proteins function are still being elucidated. In *Mtb*, secretion of ESX-1 substrates (including EsxA) is co-dependent ^65,66^. Further, in *S. aureus*, EsxA is not only necessary for secretion of other T7SS substrates ^13,19,63^, but was also co-purified with other T7SS core machinery membrane components ^20^. These studies have led to the hypotheses that T7SS substrates may be secreted as multimeric complexes and/or that WXG100 proteins such as EsxA may actually comprise part of the secretion machinery, potentially forming an extracellular or surface-associated component of the secretion apparatus ^7,54^. In this manner, recently, *Mtb* WXG100 proteins EsxEF were hypothesized to form the outer membrane T7SS channel allowing export of the toxin, CpnT ^10^. It is currently unclear whether the GBS EsxA functions as a secreted substrate or if it comprises part of the GBS T7SS machinery through association with other core components at the membrane. In support of the latter hypothesis, our data suggest that the majority of the cytotoxicity induced by the GBS T7SS is contact-dependent, as we observed minimal levels of contact-independent hCMEC cytotoxicity using transwells. Thus, GBS EsxA may also be important for the secretion of other T7SS substrates, either by chaperoning other T7SS effectors or as part of the T7SS apparatus itself; however, this requires further investigation.

In addition to its contribution to T7SS activity, EsxA is an intuitive first T7SS substrate to study as it is broadly conserved across almost all T7SS (an exception being GBS subtype IV strains, such as COH1). It has been suggested that conservation of *S. aureus* EsxA across many EssC subtypes may indicate that EsxA interacts with a conserved portion of EssC (that is common across all subtypes) instead of the EssC C-terminus, which may be specific for each subtype’s secreted effectors ^15,17^. EsxA, as well as other WXG100 proteins, commonly form ɑ-helical structures containing coiled-coil domains, and mutations of hydrophobic residues within these domains (such as the WXG motif) have been predicted to abrogate WXG100 protein interactions (either with self or with other protein partners) ^7,27,67,68^. In Tak et al, the pore-formation function of EsxEF was dependent on the WXG motif and this consequently affected secretion of the CpnT toxin ^10^. Our data herein corroborates these findings in that the WXG motif is also important for GBS EsxA-mediated cytotoxicity. Future studies will determine whether GBS EsxA-dependent cytotoxicity is specific to brain endothelium, or commonly observed in other cell types such aortic endothelium or in epithelial cells. Further, the mechanism by which EsxA or other T7SS substrates induces host cell death has been contested in previous literature and likely depends on the strain-specific secreted factors. In *Mtb*, T7SS-mediated cytotoxicity due to EsxA was shown to occur independently of pore formation, as cells died via apoptosis due to tearing of the membrane ^56^. Conversely, in *S. aureus*, EsxA was shown to inhibit or delay apoptosis ^69,70^, and in *Mtb*, export of toxin CpnT resulted in macrophage death via necroptosis ^71^. Thus, the mechanism by which GBS T7SS induces cell death requires further investigation.

Interestingly, GBS subtype I encodes two full copies of *esxA.* As these genes are 95% identical, we have annotated them as *esxA1* and *esxA2* and we show in this study that expression of just one is sufficient for maximal CJB111 cytotoxicity in brain endothelium. In addition to *esxA1* and *esxA2*, which are encoded upstream of the T7SS locus, CJB111 also encodes an orphaned WXG100 protein located elsewhere in the genome (ID870_08245) that is 86% and 84% identical to EsxA1 and EsxA2, respectively. Whether this EsxA-like protein also contributes to T7SS-mediated virulence, cytotoxicity, and pore formation is unknown.

EsxA is just one of many potential T7SS substrates in CJB111. Despite diversity in sequence and size, T7SS substrates are usually ɑ-helical in nature and often contain T7SS-associated motifs, such as WXG, LXG, YxxxD/E or the C-terminal hydrophobic pattern HxxxD/ExxhxxxH (“H” and “h” indicating highly conserved and less conserved hydrophobic residues, respectively) ^4^. Additional common substrates of the T7SS include LXG-domain containing polymorphic toxins, which encode a conserved N-terminus similar in structure to WXG100 proteins but with an extended variable C-terminal toxin domain ^72^. These toxins have been described in *S. intermedius* ^30,73^*, S. aureus* ^14,16^, and *E. faecalis* ^29^ and mediate interbacterial competition and/or host toxicity. CJB111 encodes a putative LXG toxin just downstream of the T7SS core machinery locus that has not yet been characterized. LXG toxin-encoding genes are prevalent in bacteria that comprise the human gut microbiota ^30^ and the T7SS of *E. faecalis* has been shown to be important for colonization of the murine vaginal tract ^74^. Thus, GBS LXG toxins may promote interbacterial competition of GBS with normal flora in both the gastrointestinal and female reproductive tracts and would be of interest to study in the future.

Other T7SS substrates may exist in addition to Esx proteins and LXG toxins; yet, finding conditions in which T7SS is induced *in vitro* constitutes a significant hurdle to their identification. As has been the case in studying other secretion systems, T7SS structures may only be assembled *in vivo* or when in contact with specific host factors or host or bacterial cells^7,75^. Simply identifying conditions by which to induce expression of T7SS genes *in vitro* has proven elusive ^49^ and may be species dependent. To compound these difficulties, while most Firmicutes’ T7SS loci commonly encode homologs for core T7SS machinery and WXG100 proteins, the C-terminal end of EssC as well as the downstream putative T7SS effectors (including LXG toxins) vary widely across strains of the same species. This extensive T7SS diversity has been described based on EssC C-terminal sequences in *S. aureus* (4 subtypes) ^17^, *L. monocytogenes* (7 subtypes) ^25^, and *Staphylococcus lugdenensis* (2 subtypes) ^24^. These systems exhibit little to no cross talk as in *Mtb*, ESX systems are not known to complement each other, and in *S. aureus,* expression of *essC* variants in heterologous strain backgrounds allowed EsxA secretion but not secretion of strain-specific effectors ^15^. Although only subtype I in GBS has been studied to date (present study), GBS also exhibits extensive diversity and encodes four T7SS subtypes, each associated with a unique set of effectors downstream; therefore, an EssC subtype-specific secretome likely exists across GBS strains, which will inevitably affect T7SS-dependent phenotypes. Determining whether other GBS T7SS subtypes are functional and important for virulence, as well as identifying subtype-specific effectors will be the subject of our follow-up studies.

In conclusion, this work provides the first characterization of a T7SS in GBS and is the first to demonstrate a role for a non-mycobacterial EsxA homolog in pore formation. This study builds on previous Gram-positive T7SS literature, suggesting that GBS T7SS subtype I has a role in virulence that is dependent on EsxA, and more specifically, the EsxA WXG motif. Further study of T7SS effectors may uncover previously unknown mechanisms of GBS pathogenesis and may provide insight into new therapeutic targets for Group B streptococcal disease.

## Methods

### Bioinformatic analysis of GBS T7SS

Closed genomes of *Streptococcus agalactiae* were downloaded in Geneious Prime® 2020.1.2 using the NCBI Nucleotide Blast function searching for the following terms: “Streptococcus agalactiae complete genome”, “Streptococcus agalactiae complete sequence”, or “Streptococcus agalactiae chromosome”. 136 closed genomes were downloaded in total. Protein BLAST was performed in Geneious to assess the presence of the EssC C-terminus (with the terminal 225 amino acids of CJB111 EssC used as template). Protein alignments were manually checked for true alignment to the queried sequence and strains encoding EssC exhibited a minimum Bit-score of 160, E value = 8.65e-43, Grade = 74.3%. These metrics equate to minimum of 35.9% identity/ 98.67% coverage of the 225-amino acid query. While most strains that encoded the EssC C-terminus also encoded other T7SS genes, some GBS isolates contain fragmented T7SS loci. Thus, only the EssC C-terminus sequence was assessed in this analysis. Of 136 GBS isolates, 80 encode an EssC C-terminus. A phylogenetic tree was generated in Geneious based on the EssC C-terminal sequences extracted from the above protein BLAST. Branches are transformed proportionally and are in decreasing order. T7SS subtypes were classified based on the visual branching of the tree. EsxA protein alignments were performed using the EMBL-EBI (European Molecular Biology Laboratory-European Bioinformatics Institute) ClustalW program (v.1.2.4) ^76^.

### Bacterial strains and cell lines

GBS strain CJB111, an isolate from a case of neonatal bacteremia without focus (accession: NZ_CP063198.2) ^77^ was used in this study. GBS strains were grown in Todd Hewitt Broth (THB; Research Products International, RPI) statically at 37° C. When needed, antibiotic was added to THB at final concentrations of 100 μg/mL spectinomycin. Strains containing the complementation plasmid pDCErm were grown in THB + 5 μg/mL erythromycin. All strains used in this study can be found in Table S2. The human cerebral endothelial cell line hCMEC/D3 (Millipore-Sigma; SCME-004) used in this study was grown in EndoGRO complete medium with 5% fetal bovine serum and 1 ng/mL FGF-2 (fibroblast growth factor-2).

### Cloning

Deletion mutants of *essC* (ID870_04200) and *esxA1/esxA2* (ID870_04170/ ID870_04175) genes were created as described previously using the temperature sensitive plasmid pHY304 ^78^ with slight modifications: namely, this time using a gene encoding spectinomycin resistance, *aad9*, in the knockout construct. Second crossover mutants were screened for erythromycin sensitivity and spectinomycin resistance. Vector controls and complemented mutants were generated as previously described using overexpression plasmid pDCErm ^78^. Primers used in this study can be found in Table S3.

### RNA purification and qRT-PCR

qRT-PCR analysis of bacterial gene expression was performed as described previously ^78^. To assess gene expression in CJB111, Δ*essC* mutant, and *essC* complement, strains were grown to mid-log (OD_600_ = 0.4 - 0.6) and RNA was purified using the Machery-Nagel Nucleospin kit (catalog# 740955.250) according to manufacturer instructions with the addition of three bead beating steps (30 sec × 3, with one minute rest on ice between each) following the addition of RA1 buffer + β-mercaptoethanol. Purified RNA was treated with the turbo DNAse kit (Invitrogen, catalog# AM1907) according to manufacturer instructions. cDNA was synthesized using the SuperScript cDNA synthesis kit (QuantaBio, catalog# 95047-500), per manufacturer instructions. cDNA was diluted 1:150 to further reduce bacterial DNA contamination and qPCR was performed using PerfeCTa SYBR Green (QuantaBio, catalog# 95072-05K) and BioRad CFX96 Real-Time System, C1000 Touch Thermocycler. qRT-PCR primers used in this study can be found in Table S3.

### Murine model of hematogenous meningitis

Contribution of T7SS to GBS virulence was assessed using a model of hematogenous meningitis as described previously ^78^. Male 8-week old CD1 mice (Charles River) were tail vein injected with 2 - 3 × 10^7^ CFU of CJB111 or an isogenic T7SS mutant. Mice were sacrificed upon exhibition of neurological symptoms such as paralysis or moribundity in accordance with our IACUC protocol # 00316 (University of Colorado Anschutz Medical Campus). Upon mouse death, brain, heart and blood were collected. Tissue was homogenized and samples were serially diluted and plated on THA for CFU enumeration. Bacterial counts were normalized to the tissue weight.

### ELISA

KC protein in homogenized tissues was quantified using R&D systems ELISA kits (catalog # DY453). KC protein detected was normalized to tissue weight and reported as KC protein (pg) per mg of tissue.

### Cell based assays: Adherence, Invasion, Intracellular survival, and LDH release

hCMECs were passaged, seeded at 150,000 cells/well into rat tail collagenized 24-well plates (Corning, catalog# 3524), and grown into a confluent monolayer overnight in EndoGRO complete medium. Confluent cell monolayers were then washed with PBS and media was replaced (0.4 mL media per well). For cell-based assays, GBS was sub-cultured from overnight cultures (1:10) and grown to mid-log. Bacteria were pelleted and normalized to an OD_600_ value predetermined to yield 1 × 10^8^ CFU in PBS. Bacteria were serially diluted in PBS and added to the cell monolayers in 24-well plates.

Adherence, invasion, and intracellular assays were performed as previously described ^78^. Briefly, for adherence assays, GBS was added to hCMEC monolayers at an MOI of 1 and incubated for 30 minutes. For invasion assays, GBS was added to hCMEC monolayers at an MOI of 1, incubated two hrs, washed three times with PBS, and then incubated in media containing penicillin and gentamycin for two hrs to kill any extracellular bacteria. In both these assays, at the final timepoint, cells were washed with PBS, trypsinized five minutes at 37° C, and lysed using 0.025% Triton-X-100 in PBS. After mixing the lysate well by pipetting, CFU were quantified by serial dilution of the lysate and plating on THA. Percent adherence or invasion was calculated by taking the quotient of inoculum and the CFU quantified at the end of the assay. Intracellular survival assays were performed identically to the invasion assay, except cells were incubated for 12 hours instead of 2 hours following the addition of antibiotic-containing medium.

For LDH release assays, bacteria were added to hCMEC monolayers as described above in EndoGRO complete medium at an MOI of 10 and allowed to incubate for 4-5 hrs. LDH release was measured according to manufacturer instructions (Pierce, Thermo Fisher, catalog # 88953).

Transwell assays were performed in tissue culture-treated 24-well polystyrene plates containing 6.5mm, polycarbonate transwell inserts with 0.4μM porous mebranes (Corning, catalog# 3413). hCMEC were seeded into collagenized wells in the bottom compartment and grown overnight in EndoGRO complete medium as described above. On the day of the assay, cells were washed with PBS, fresh medium was added, and transwells were inserted and equilibrated with EndoGRO complete medium. GBS was added to the transwell bucket at an MOI of 10, and the plate was incubated for 24 hours. Supernatant from the hCMEC lower compartment was plated at the end of the assay to ensure lack of bacterial contamination.

### EsxA1 protein purification

Purification of CJB111 EsxA1 was performed as recently described ^10^ with modifications described here. The *esxA1* gene of CJB111 (ID870_04170) was cloned into expression vector pML3339 ^79^ and expressed in *Escherichia coli* BL21(DE3) using 1L of ZYP5052 autoinduction medium ^80^ + carbenicillin (100 μg/ml). Cultures were grown at 37°C, shaking (200 rpm) for 24 hours and the *E. coli* pellet was resuspended in lysis buffer (50 mM HEPES, 150 mM NaCl pH 7.5 + 1 mM PMSF, + 1 mg/ml lysozyme, + 3 μl benzonase (Novagen, Merck Millipore 70746-3), + 1 Roche complete protease inhibitor tablet), sonicated 30 seconds on /off for 20 minutes on ice, and spun at 14,000 × g to pellet debris and to collect the soluble fraction. The soluble fraction was run over packed and equilibrated nickel resin (Thermo Fisher, HisPur™ Ni-NTA Resin, catalog# 88222), washed with 25 mM HEPES 150 mM NaCl + 20 mM imidazole (pH 7.0) and eluted in 25 mM HEPES 150 mM NaCl + 300 mM imidazole (pH 7.0) to obtain clean 6xHIS-MBP-EsxA (maltose binding protein; fusion protein is~55 kDa). The sample was dialyzed (3.5 kDa MWCO, Spectrum™ Spectra/Por™, catalog# 086705B) to remove imidazole, cleaved with 6xHIS-TEV protease overnight at 4°C, and then run over a nickel column to bind 6xHIS-MBP and 6xHIS-TEV as described above. EsxA1 was collected in the flowthrough, run over amylose resin (NEB, catalog# E8021L) twice to bind any remaining MBP contaminants, and the flowthrough was collected. The final EsxA1 product was concentrated using 3 kDa amicon centrifugal filters (Millipore, catalog# UFC900324) and verified by Coomassie, where it exhibited a clear monomeric band at 10 kDa in addition to putative dimers, trimers, and higher order oligomers.

EsxA sample purity was confirmed using tryptophan fluorescence (NanoTemper Tycho). Pure MBP was used as a negative control (sample impurity control) and EsxEF ^10^ was used as a positive control. To further confirm the purity of the sample, purified EsxA (including monomers and oligomers) was incubated with either water or guanidine hydrochloride (6 M) for 30 minutes at room temperature, run on a native gel (12% Mini-PROTEAN® TGX™ Precast Protein Gels, BioRad), and transferred to a membrane using BioRad’s Trans-Blot Turbo Transfer System (high molecular weight settings). Membranes were washed three times in TBST and blocked in LI-COR’s Intercept® Blocking Buffer (catalog# 927-60001) for one hour at room temperature. Membranes were probed with an ammonium sulfate cut of anti-EsxA1 rabbit antiserum (30 μg/ml; GenScript) or murine anti-MBP monoclonal antibody (1:10,000; NEB; catalog# E8032S) in the above LI-COR blocking buffer, overnight at 4° C. Following washes in TBST, membranes were incubated with IRDye 680RD goat anti-rabbit or goat anti-mouse IgG (H + L) secondary antibodies from LI-COR (1:10,000 dilution; 1 hour, room temperature; catalog#s 926-68071 and 926-68070, respectively). Following washes in TBST and water, western blots were imaged using the LI-COR Odyssey.

### Lipid bilayers

Pore forming activity was assessed using lipid bilayers as described previously ^10^ using EsxA1 protein that had been purified that day. Briefly, 100% DphpC (1,2-diphytanoyl-sn-glycerol-3-phosphocholine, 4ME 16:0 PC) lipid bilayers were used in 25 mM sodium phosphate, 1M KCl at pH 7.4 or pH 4.0. Purified GBS EsxA1 (5 μg) was added to cis / trans side and nine membranes were assessed for pore-forming activity. 46 insertions were observed in total. The insertion profile did not exhibit a gaussian distribution or trend towards a particular conductance value. At pH 4.0, 132 insertions were observed in total. The insertion profile at pH 4.0 exhibited a more uniform distribution and, once channels were formed, the overall conductance was higher than those at pH 7.4. Data was analyzed using a custom algorithm in IGOR Pro.

### Transmission Electron Microscopy of negatively-stained EsxA1

Negative stain of EsxA1 was performed as described previously ^10^. Recombinant EsxA1 was prepared to a concentration of approximately 370 μg/mL in 25mM sodium phosphate buffer pH 4.0, blotted on glow-discharged grids (continuous carbon), washed twice with milli-q water, and then stained with 1% uranyl formate for 2 minutes. All grids were prepared within 16-48 hours of EsxA1 purification. Micrographs were collected on a 120kV ThermoFisher Talos L120C transmission electron microscope using 45,000X magnification and a fixed defocus of −2.24 μm. Particle picking was performed using EMAN2.2 Swarm picking with a particle size of 100 and box size of 150 at 3.19 Å/ pixel. Particle quality was manually inspected and aggregates were removed. Reference-free 2D class averages were generated in EMAN2.2 from a total of 12,727 particles over 43 CTF-corrected micrographs. Micrograph quality / CTF was confirmed manually. The low-pass filtered (20 Å) particle set was subjected to four iterations of class averaging. Each iteration displayed similar results.

## Supporting information

Table S1--GBS strains by T7SS subtype

Table S2--Strains used in this study

Table S3--Primers used in this study

## Acknowledgments

We thank Jennifer Bourne and Eduardo Romero Camacho of the Electron Microscopy Core and the Cryo-EM Structural Biology Shared Resource Facility (University of Colorado Anschutz Medical Campus), respectively, for assistance with negative stain and imaging of EsxA1 protein. We also thank Terje Dokland (University of Alabama at Birmingham) for his review of our negative strain/transmission electron microscopy data. Funding for this work was provided by NIH/NIAID F32 AI143203 to BLS, the Coordenação de Aperfeiçoamento de Pessoal de Nível Superior - Brasil (CAPES) - Finance Code 001 to JCM, and NIH/NINDS R01 NS116716 and NIH/NIAID R01 AI153332 to KSD.

## Author Contributions

BLS and KSD conceived the project. BLS, UT, and KSD designed the experiments and BLS, UT, and JCM acquired the data. PEN and MN provided resources and BLS and KSD wrote the manuscript. All authors revised the final manuscript and approved of its submission.

