## Supplemental Figures 1-4 for "A type VII secretion system in Group B *Streptococcus* mediates cytotoxicity and virulence"

**Running title**: GBS T7SS mediates virulence

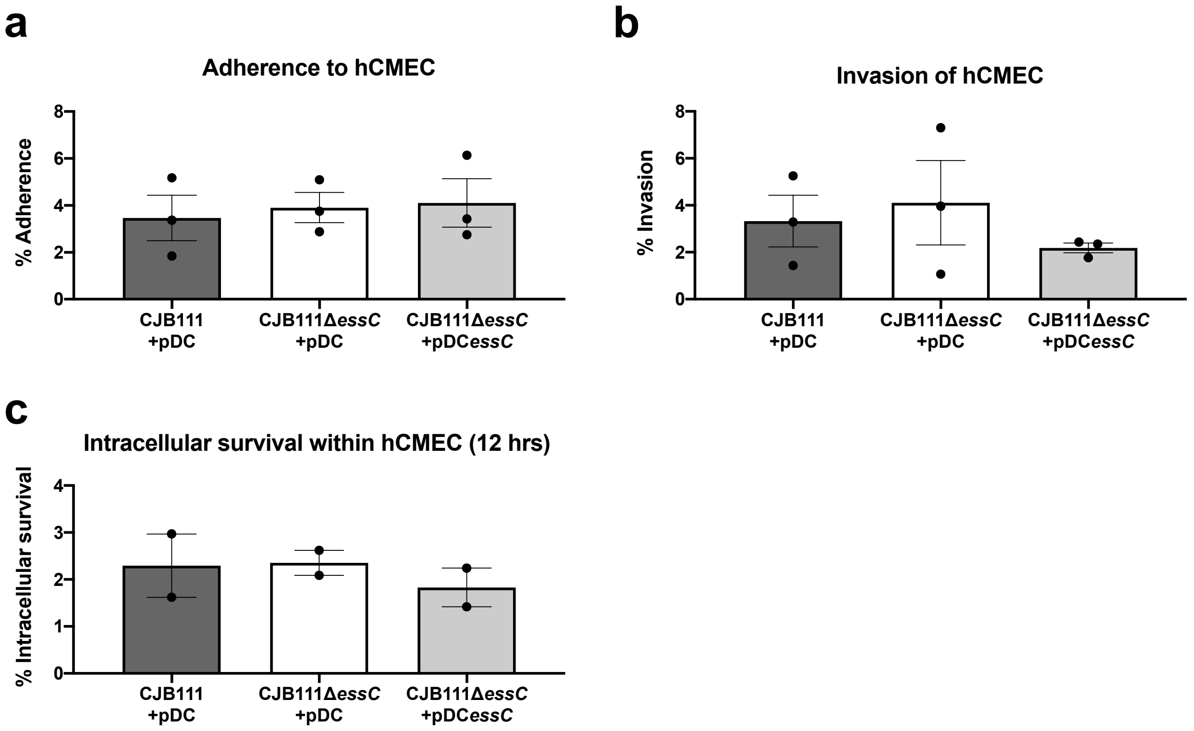


**Figure S1. Deletion of *essC* does not affect GBS interaction with host cells, *in vitro.***

CJB111 and the Δ*essC* mutant were further evaluated for **a)** adherence to (n = 3), **b)** invasion of (n = 3), or **c)** intracellular survival (12 hrs; n = 2) in human cerebral microvascular endothelial cells (hCMEC). Data represent percent CFU recovered of the initial inoculum and were performed in technical duplicates or triplicates.

**a**

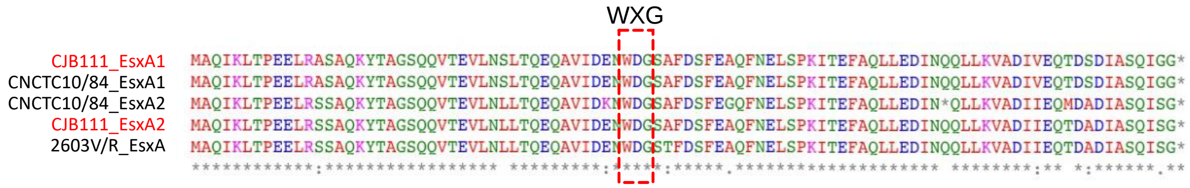


**b**

| | CJB111_EsxA1 | CNCTC10/84_EsxA1 | CNCTC10/84_EsxA2 | CJB111_EsxA2 | 2603V/R_EsxA |
|---|---|---|---|---|---|
| CJB111_EsxA1 | | 100 | 76.84 | 94.79 | 93.75 |
| CNCTC10/84_EsxA1 | 100 | | 76.84 | 94.79 | 93.75 |
| CNCTC10/84_EsxA2 | 76.84 | 76.84 | | 77.89 | 76.84 |
| CJB111_EsxA2 | 94.79 | 94.79 | 77.89 | | 98.96 |
| 2603V/R_EsxA | 93.75 | 93.75 | 76.84 | 98.98 | |

**Figure S2. Canonical T7SS substrate EsxA is conserved across GBS T7SS subtypes I - III.**

**a)** ClustalW alignments and **b)** percent identity matrix of EsxA amino acid sequences from GBS T7SS subtypes I-III. Subtype I encodes EsxA1 and EsxA2 and is represented by CJB111 (accession: NZ_CP063198.2). Subtype II encodes EsxA2 and is represented by 2603V/R (accession: NC_004116.1). Subtype III encodes EsxA1 and a truncated EsxA2 and is represented by CNCTC 10/84 (accession: NZ_CP006910.1). In the above matrix, the purple shading corresponds to the level of identity between two strains (on a spectrum of 0 to 100% identity), with darker shading indicative of higher percent identity.

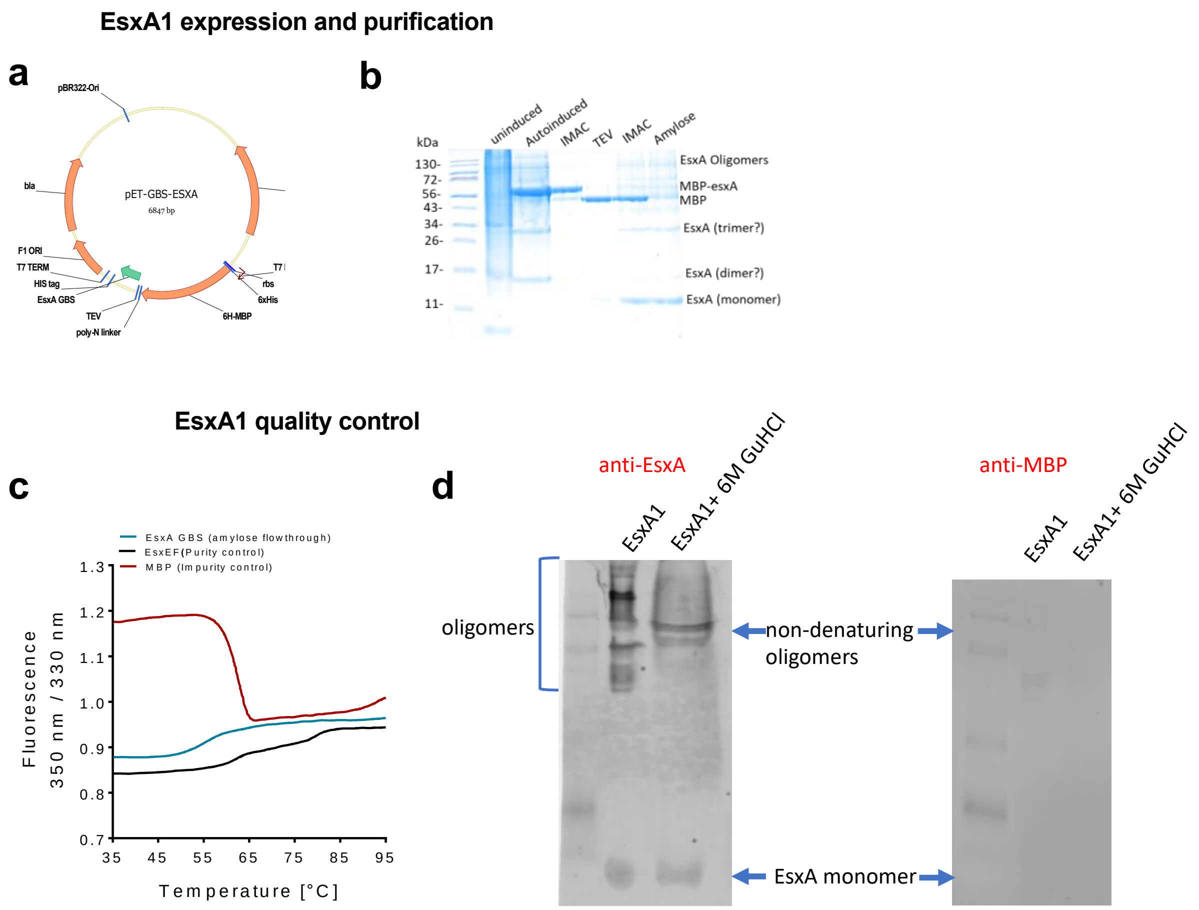


**Figure S3. EsxA1 expression, purification, and quality control.**

**a)** Plasmid map of CJB111 *esxA1* cloned into pET vector pML3339 from Michael Niederweis. EsxA1 expressed from this vector is His-tagged and MBP-tagged to facilitate nickel affinity and amylose affinity column purification.

**b)** SDS-PAGE gel of EsxA1 during purification: un-induced BL21 culture, auto-induced BL21 culture, post-nickel affinity column (IMAC), post-TEV cleavage/dialysis; post amylose column to remove cleaved MBP, final EsxA1. Final EsxA1 product shows a ~10.6 kDa monomer as well as higher order oligomers.

**c)** Quality control of the purified EsxA1 by Tycho (NanoTemper Technologies). Maltose binding protein was run as a negative control and mycobacterial EsxEF was run as a positive control.

**d)** Native-PAGE indicating that most EsxA1 oligomers resolve to the monomeric state upon treatment of protein with 6M guanidine HCl for 30 minutes at room temperature. EsxA1 bands stain with anti-EsxA1 rabbit antiserum but not with anti-MBP antibody.

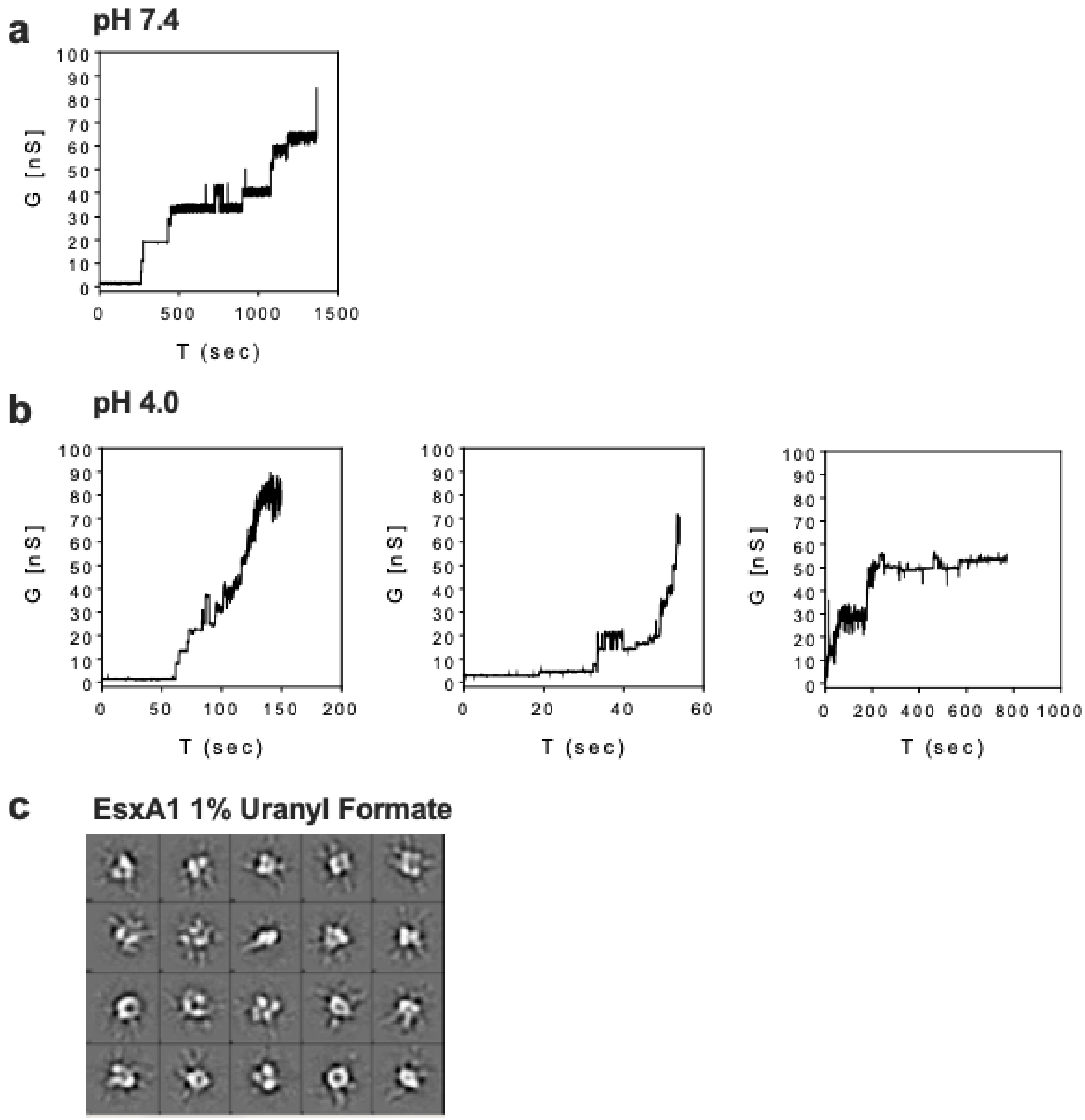


**Figure S4. Complete set of EsxA1 lipid bilayer traces and 2D class averages.**

Additional current traces of recombinant CJB111 EsxA1 pore formation in DphpC lipid bilayers at **a)** pH 7.4 and **b)** pH 4.0 in 25 mM sodium phosphate 1M KCl. **c)** Additional reference-free 2D class averages of negatively stained EsxA1 imaged by transmission electron microscopy.
